# Super-resolution imaging of neuronal structure with structured illumination microscopy

**DOI:** 10.1101/2023.05.26.542523

**Authors:** Tristan C. Paul, Karl A. Johnson, Guy M. Hagen

## Abstract

Super-resolution structured illumination microscopy (SR-SIM) is a method in optical fluorescence microscopy which is suitable for imaging a wide variety of cells and tissues in biological and biomedical research. Typically, SIM methods use high spatial frequency illumination patterns generated by laser interference. This approach provides high resolution but is limited to thin samples such as cultured cells. Using a different strategy for processing the raw data and coarser illumination patterns, we imaged through a 150 µm thick coronal section of a mouse brain expressing GFP in a subset of neurons. The resolution reached 144 nm, an improvement of 1.7 fold beyond conventional widefield imaging.

## Introduction

Recently developed methods for surpassing the diffraction limit in optical fluorescence microscopy include stimulated emission depletion microscopy (STED) [1], stochastic optical reconstruction microscopy (STORM) [2], photoactivated localization microscopy (PALM) [3], super-resolution optical fluctuation imaging (SOFI) [4], and structured illumination microscopy (SIM) [5,6]. These methods have had large impacts in many fields, with super-resolution microscopy previously being used to image the mouse brain with STED [7], STORM [8], and SIM approaches [9].

SIM is a method in which sets of images are acquired with shifting illumination patterns. Subsequent processing of these image sets yields results with optical sectioning, resolution beyond the diffraction limit (super-resolution), or both [5,6,10–13]. Since its emergence over two decades ago [14], SIM has matured as an imaging technique, with multiple proposed methods for generation of the structured illumination patterns [11–24] as well as for processing of the image data [6,13,25–29]. Compared to other super-resolution techniques, the speed, high signal-to-noise ratio, and low excitation light intensities characteristic to SIM make it a good choice for imaging of a variety of samples in three dimensions.

The imaging method we used, maximum *a posteriori* probability SIM (MAP-SIM) uses a Bayesian framework to reconstruct super-resolution SIM images [25,28,30]. This method has advantages including flexibility in the range of SIM illumination patterns which can be used, in our case the ability to use patterns with lower spatial frequencies. This in turn allows imaging deeper into samples in which scattering degrades the high spatial frequency patterns which are more commonly used in SIM. When the SIM pattern is out of focus, it blurs rapidly with increasing depth, producing a high intensity of out of focus light. This results in reduced pattern contrast in the acquired images. Because of this, traditional super-resolution SIM methods are typically limited to an imaging depth of 10-20 µm [31,32]. Here we used MAP-SIM to image a fixed, optically cleared, ∼150 µm thick mouse brain coronal slice expressing a neuronal GFP marker while achieving a lateral resolution of 144 nm.

To overcome the challenges of imaging deeper into brain tissues, SIM has previously been combined with adaptive optics for in vivo studies [33,34]. These methods offered impressive results, but they do require additional optical devices and expertise. Here we used a simpler and more economical approach with a non-laser light source and open-source software for SIM [35].

## Methods

The sample used for this work was an optically cleared, green fluorescent protein (GFP)-labeled coronal mouse brain slice. The slice was approximately 150 µm thick and was obtained from SUNJin Lab (Hsinchu City, Taiwan).

We used a home-built SIM set-up based on the same design as described previously [15,25,30,36]. Briefly, the SIM system is based on an IX83 microscope (Olympus, Tokyo, Japan). Illumination was provided by a liquid light guide-coupled Spectra-X light source (Lumencor, Beaverton, Oregon) using the cyan channel, which has an emission maximum of 470 nm. The illumination was collimated by an achromatic 50 mm focal length lens (Thor labs, Newton, New Jersey) and polarized with a linear polarizer (Edmund Optics, Barrington, New Jersey) before passing into a polarized beam splitter (PBS) cube (Thor Labs) and illuminating a Liquid Crystal on Silicon (LOCS) microdisplay (SXGA-3DM, 13.6 μm pixel pitch, Forth Dimension Displays, Dalgety Bay, Scotland). The output of the microdisplay again passed through the PBS and was imaged into the microscope using a 180 mm focal length lens (SWTLU-C, Olympus). For this work, two objectives were used, UPLAPO 10×/0.4 NA water immersion, and UPLSAPO 100×/1.4 NA oil immersion (Olympus). Emitted fluorescent light was filtered (GFP filter set with dichroic T495lpxr and ET525/50 emission filter, Chroma, Bellows Falls, Vermont) and then imaged with a sCMOS camera (Zyla 4.2+, Andor). Sample movements and focusing were controlled by a XY, piezo Z stage (Applied Scientific Instrumentation, Eugene, Oregon).

The microdisplay was used to produce the SIM patterns and was controlled by the software supplied with the device (MetroCon, Forth Dimension Displays). For these experiments, the pattern consisted of two columns of on-pixels followed by 10 rows of off pixels repeated across the microdisplay. This pattern was shifted horizontally by one pixel after each image was acquired. Twelve total images were acquired per z-slice so that the sum of all illumination masks resulted in homogenous illumination across the sample. Figure 1 shows a simplified diagram of the SIM optical system and a connection diagram illustrating how the microdisplay system is synchronized with the camera using IQ software (Andor) and a digital input/output computer card (DDA06/16, Measurement Computing, Concord, New Hampshire).

**Figure 1:**
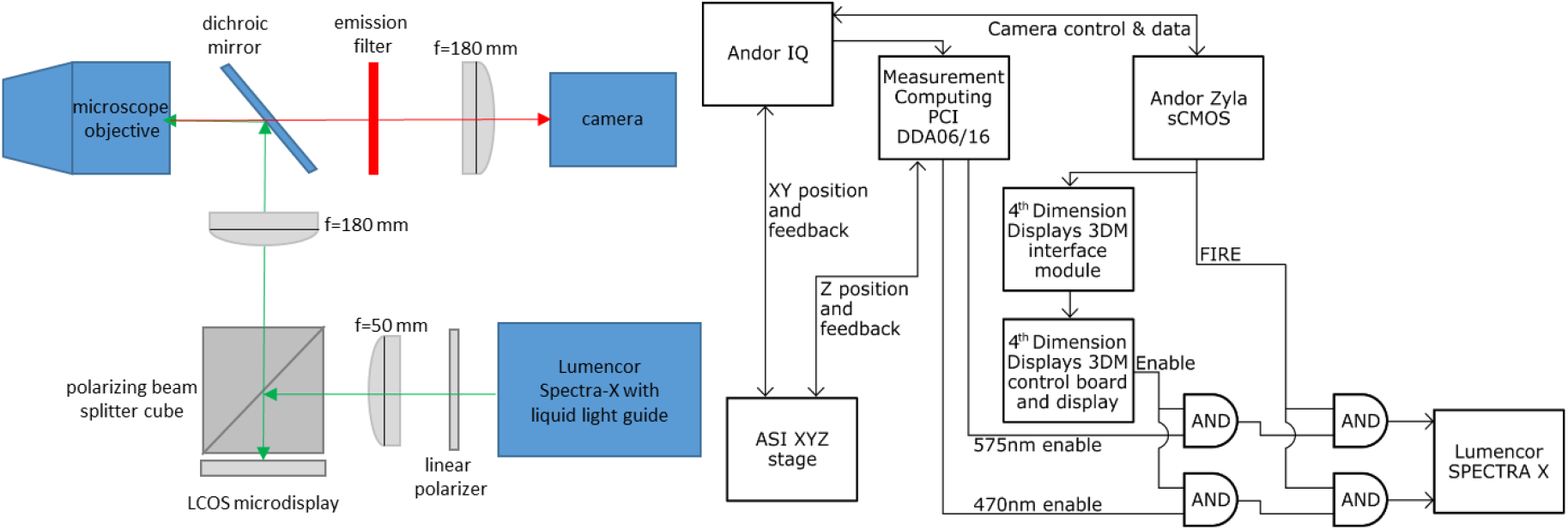
(left) simplified optical diagram; (right) connection diagram. Shown is the connection setup for two-wavelength acquisition, in this study only 470 nm illumination was used.

## Data Analysis

### Optical Sectioning SIM (OS-SIM)

Several data processing methods are possible for generating optically sectioned images from SIM data (OS-SIM) [15,37]. The most commonly used implementation of this technique was introduced in 1997 by Neil et al [14]. Their method works by projecting a line illumination pattern onto a sample, followed by acquisition of a set of three images with the pattern shifted by relative spatial phases 0, 2π/3, and 4π/3. An optically sectioned image can be recovered computationally as

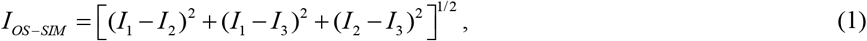

where *I*_*OS-SIM*_ is an optically sectioned image, and *I*_*1*_, *I*_*2*_, and *I*_*3*_ are the three images acquired with different pattern positions. If the sum of the individual SIM patterns results in homogeneous illumination, as is the case in our setup, a widefield (WF) image can be recovered from the SIM data by taking the average of all images:

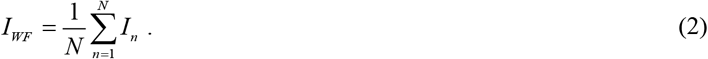

This is the approach we used throughout this study to generate conventional widefield images.

Instead of using Eq. 1, in this study we used a method originally shown by Neil et al [14] and later elaborated upon [15,37],

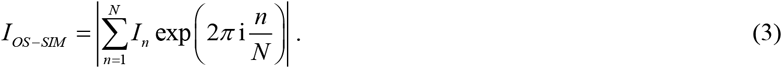

We previously showed that this processing method offers results with better optical sectioning than the method of Eq. 1 [15].

### SIM with maximum a posteriori probability estimation

MAP-SIM has been described previously [25]. In this case, the imaging process can be described as

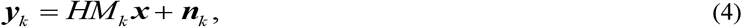

where *M*_*k*_ represents the *k*-th illumination pattern, ***y***_*k*_ denotes a low-resolution (LR) image acquired using the *k*-th illumination pattern, ***x*** is an unknown, high-resolution (HR) image, and ***n***_*k*_ is additive noise. *H* is a matrix describing the convolution between the HR image and the point spread function (PSF) of the system. The position of the illumination patterns in the camera images were determined using a calibrated camera according to our previous work [15]. We model the PSF as an Airy disk, which in Fourier space leads to an optical transfer function (OTF) of the form [38]

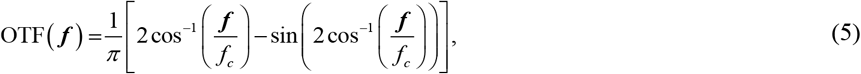

where ***f*** is the spatial frequency. We estimate the cut-off frequency *f*_*c*_ by calculating the radial average of the power spectral density (PSD) of a widefield image of 100 nm fluorescent beads.

Using a Bayesian approach [25,26,28,39–41], high-resolution image estimation can be expressed as a minimization of a cost function according to

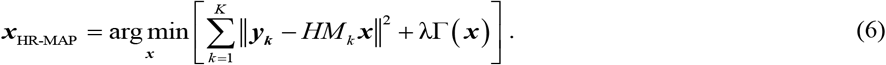

The cost function in Eq. 6 consists of two terms. The first term describes the mean square error between the estimated HR image and the observed LR images. The second term is a regularization term. To ensure positivity and promote a smoothness condition, we rely on quadratic regularization composed of finite difference approximations of the first order derivative at each pixel location [42]

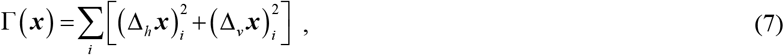

where Δ_*h*_ and Δ_*v*_ are the finite difference operators along the horizontal and vertical direction of an image. The contribution of Г(***x***) is controlled by the parameter λ, a small positive constant defining the strength of the regularization (typically λ = 0.01). We solve Eq. 6 using gradient descent methods.

### Spectral merging

MAP estimation of high-resolution images obtained with structured illumination enables reconstruction of high-resolution images (HR-MAP) with details unresolvable in a widefield microscope. However, MAP estimation as described above does not suppress out of focus light. On the other hand, the processing method according to equation 3 used in optical sectioning SIM [14,15] provides images (LR-HOM) with optical sectioning. Noting that the unwanted out of focus light is dominant at low spatial frequencies, we merge the LR-HOM and HR-MAP images in the frequency domain to obtain the final HR image (MAP-SIM). Frequency domain Gaussian low pass filtering is applied to the LR-HOM image and a complementary high pass filter is applied to the HR-MAP image. We use a weighting scheme which can be described by

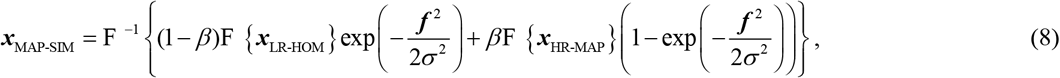

where F, F ^−1^ denotes the Fourier transform operator and its inverse, respectively, ***f*** is the spatial frequency, *σ* is the standard deviation of the Gaussian filter, and *β* is a weighting coefficient. Usually, we set *β* = 0.85. We typically use a standard incoherent apodizing function to shape the final MAP-SIM spectrum before the final inverse FFT.

## Results

To acquire an overview of the slice with SIM methods, we first imaged using a 10x/0.4 NA water immersion objective. 60 image positions were acquired with a 20 percent overlap between each position and with 12 z-planes. Image stitching was accomplished using our lab’s methods and an ImageJ plugin [43] as shown in [36]. The composite image of the slice is shown in figure 2 and is color-coded based on depth using the isolum color table [44]. This image was acquired in 5 min 30 seconds, with an additional ∼15 minutes required for OS-SIM processing according to Eq. 3.

**Figure 2:**
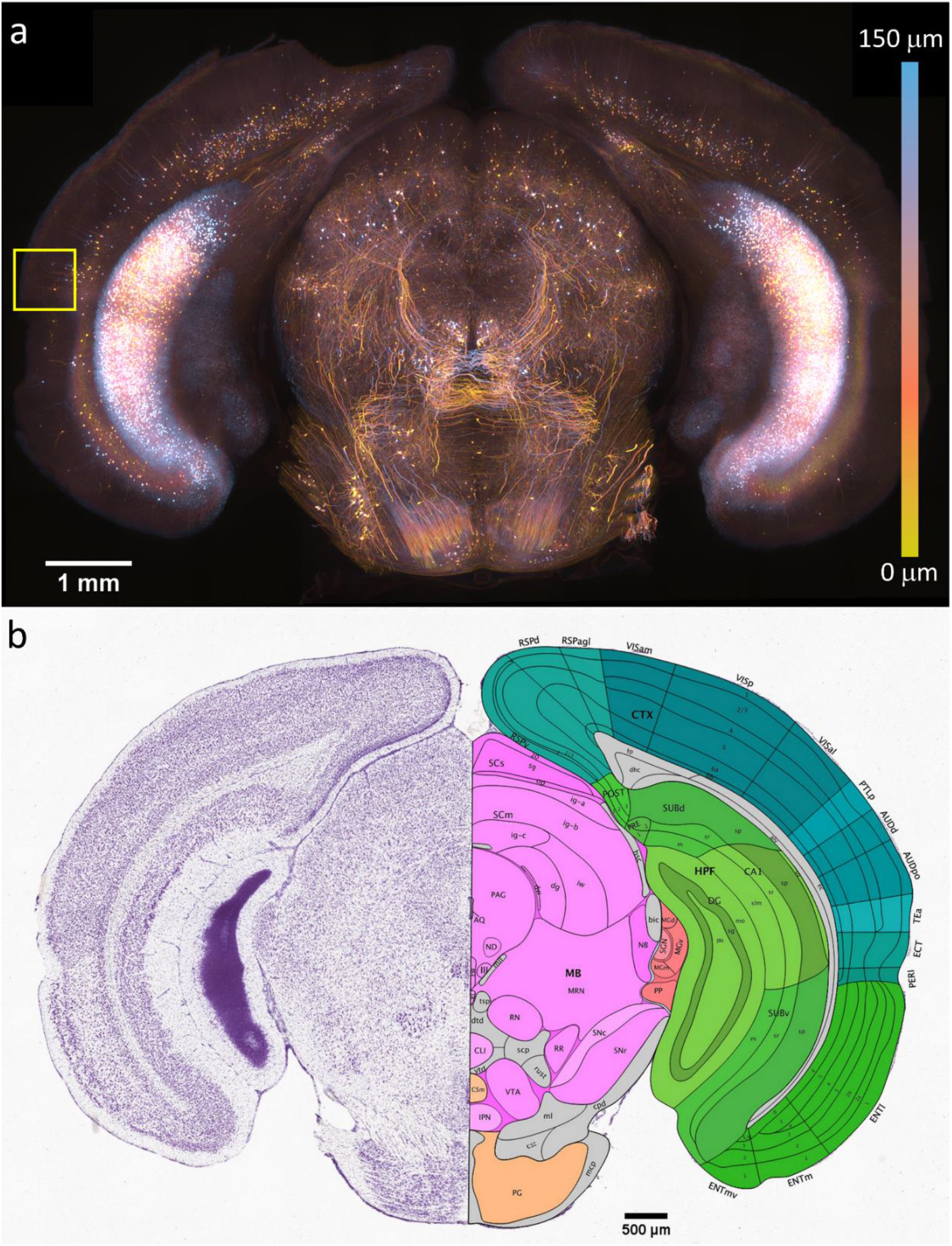
(a), Overview SIM image. The yellow box indicates the temporal association area where neurons were imaged with super-resolution MAP-SIM. (b), Allen brain atlas slice 92 of 132.

This slice was matched to Paxinos and Franklin’s mouse brain atlas [45] to identify which section of the brain was being imaged. This slice was visually matched with figure 64. We further matched our sample to slice 92 of 132 in the Allen brain atlas [46]. Then second order polynomial fits were made for both the horizontal and vertical directions using the edges and the central aqueduct as reference points. This allows any point on this brain slice, recorded from the microscope stage coordinates, to be translated into the atlas’s coordinates. Anytime the sample was moved, the location of the central aqueduct was recorded and used as the origin to account for offsets. This method placed the neuron shown in figure 3 in the temporal association area (TeA) of the mouse brain isocortex as indicated in figure 2.

**Figure 3:**
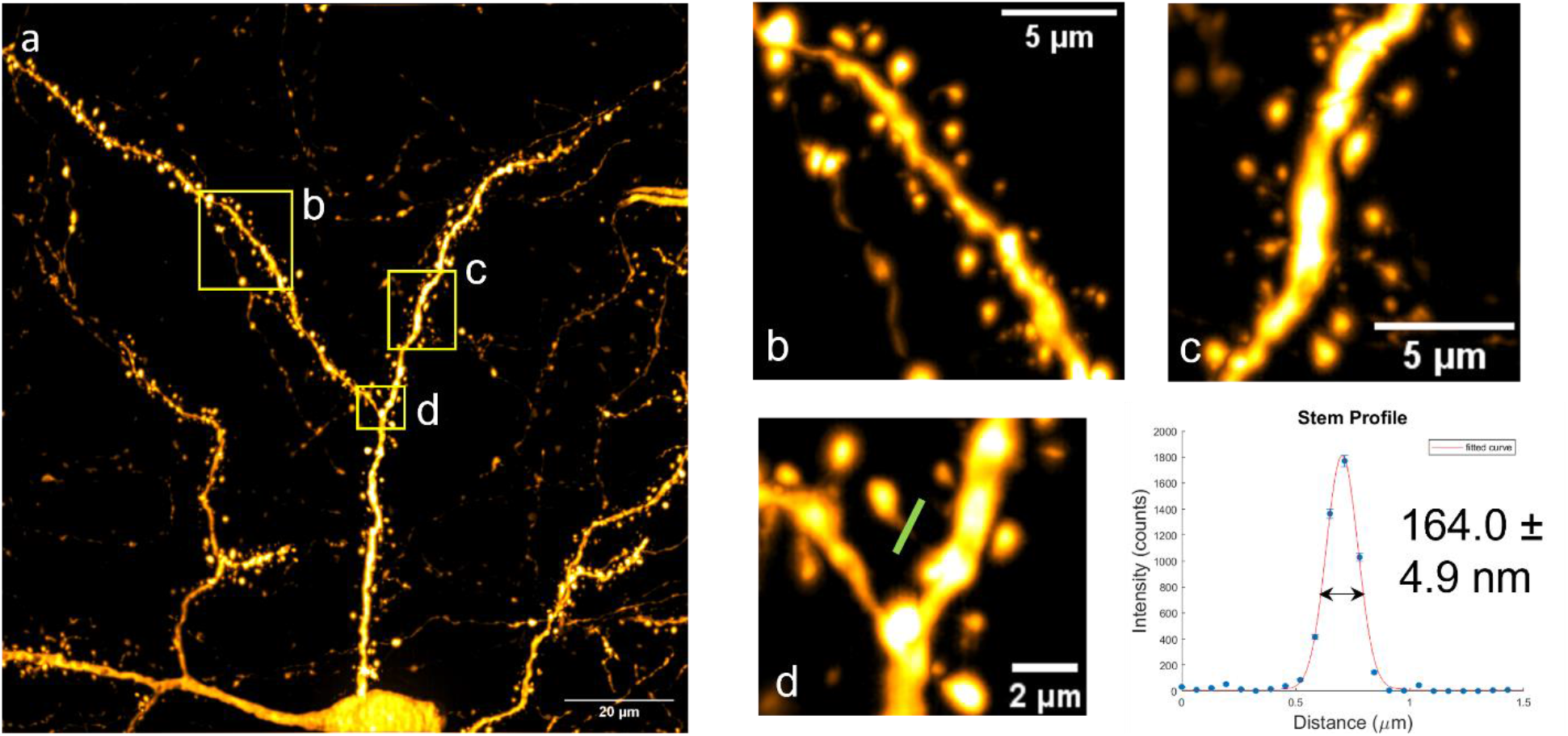
(a), neuron imaged at a depth of 41 µm to 66 µm using a 100x / 1.4 NA oil immersion objective. (b-c) show zoomed in views of the selected areas indicated in a by yellow boxes. The width of the spine neck, selected in (d) was fit to a Gaussian function, FWHM 164.0 ± 4.9 nm.

### 2.2 Imaging Deep Neurons

To demonstrate MAP-SIM’s ability to image deeper into the sample than traditional SR-SIM, a TeA neuron 41-66 µm deep was imaged. Depth was measured using the closed-loop piezo stage. A 100×/1.4 NA oil immersion objective was used with an acquisition time of 300 ms per SIM phase. This image is shown in Figure 3. To further test the resolution of deep sample MAP-SIM, the profile of a dendric spine’s neck was measured. This is shown in Fig. 3d and 3e. The profile was fit in MatLab using a Gaussian function weighted by the square root of the counts with nonlinear least squares methods. The full width at half max (FWHM) was determined to be 164.0 ± 4.9 nm.

A comparison of widefield, basic OS-SIM, and MAP-SIM is shown in Figure 4 and further analyzed in table 1. As is evident in the figure, widefield has the largest background due to out of focus light, with basic SIM providing optical sectioning and MAP-SIM providing both optical sectioning and super-resolution. The imaging depth of 41-66 µm exceeds the depth limit of traditional SIM by up to ∼3 fold. To determine the resolution, we calculated the power spectral density as previously described [47] We found that the WF image had a resolution of 247.6 nm while the MAP-SIM image had a resolution of 143.6 nm, an improvement of ∼1.7 fold. These results are summarized in Table 1.

**Table 1:** SNR and resolution measurements

|  | SNR (dB) | Resolution (nm) |
| --- | --- | --- |
| **Widefield** | **43.87** | **247.6** |
| **Basic SIM** | **29.33** | **251.6** |
| **MAP-SIM** | **39.27** | **143.6** |

**Figure 4:**
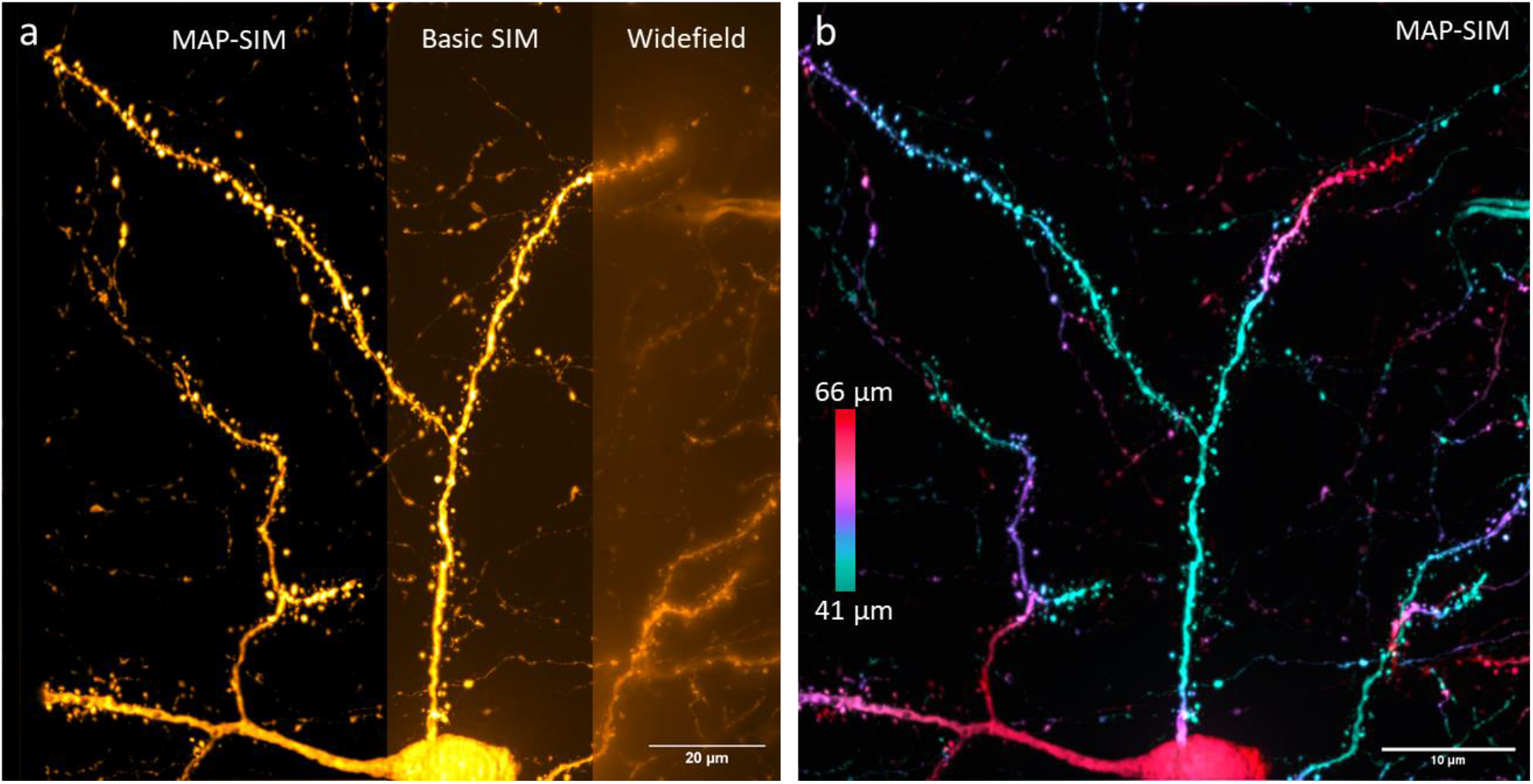
(a), TeA neuron shown in widefield, basic SIM, and MAP-SIM. (b), MAP-SIM image color coded by depth.

Typical SIM uses high frequency patterns to maximize the obtainable resolution but has limited imaging depth due to scattering and generation of large amounts of background fluorescence. The pattern used here uses a lower spatial frequency with thicker lines of illumination to penetrate deeper into the mouse brain while maintaining pattern integrity. A comparison of (cropped) images acquired using a high frequency pattern (one out of five microdisplay pixels activated) and our lower frequency pattern (two out of ten microdisplay pixels activated) of a neuron at a depth of 51 µm is shown in Figure 5. The high frequency pattern (Figure 5a) has a calculated resolution of 152 nm compared to 157 nm for the lower frequency pattern (Figure 5b). This minimal decrease in resolution is made up for by decreased imaging time and increased signal strength. Both Figure 5a and 5b have been scaled equally in brightness to highlight the increased signal strength of the lower frequency pattern despite a 50% longer exposure time used in Figure 5a.

**Figure 5:**
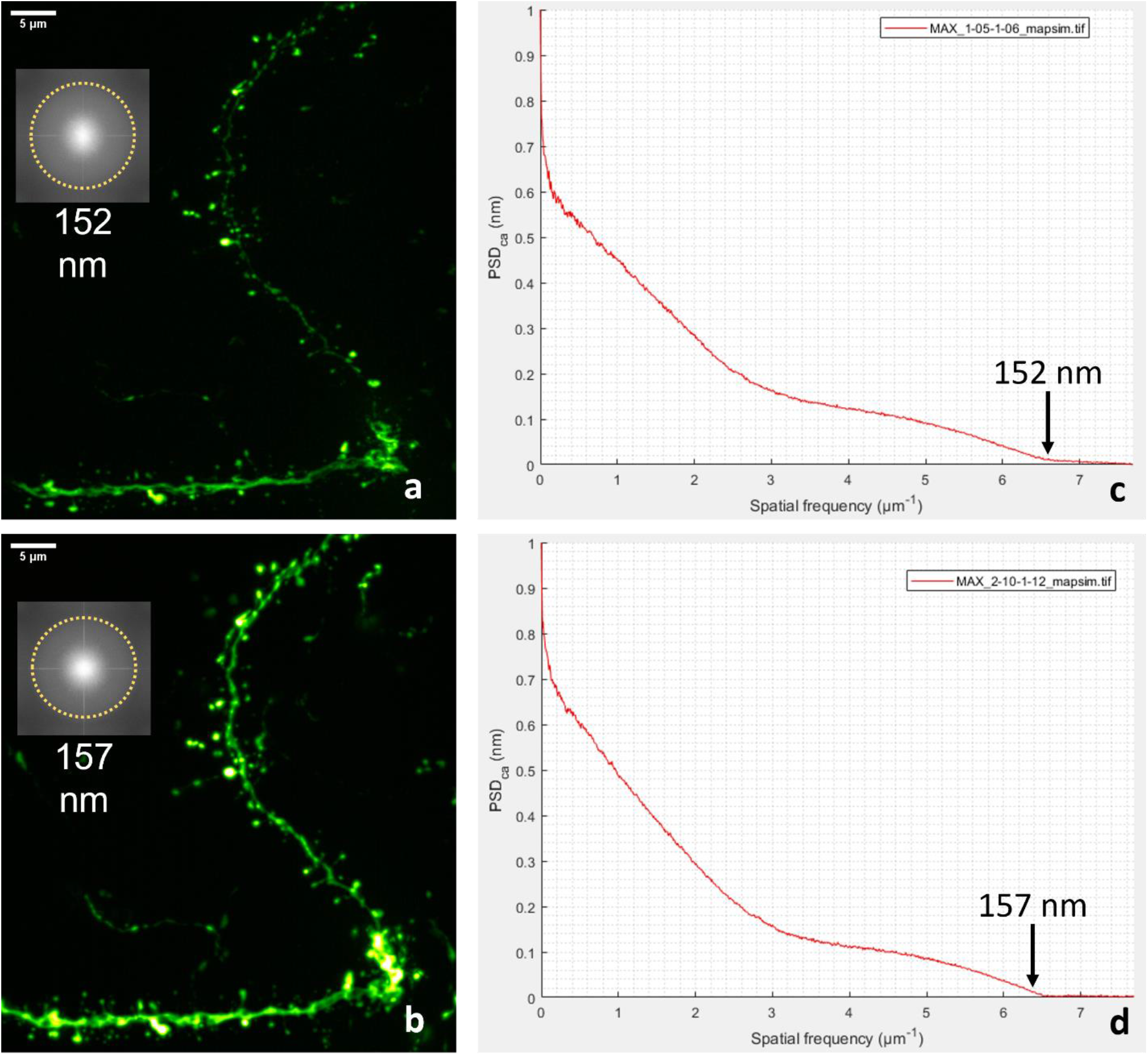
Neuron imaged using a high frequency (1 pixel on, 5 off) pattern (a) and a low frequency (2 pixels on, 10 off) pattern (b) as well as graphs of their circularly averaged power spectral density (c,d). Insets in images (a,b) are the corresponding FFTs with yellow dotted circles indicating the region of support.

## Conclusion

By combining a structured illumination microscope with a large field of view and an image reconstruction method based on Bayesian statistics, we demonstrated synapse-resolving mesoscale volumetric imaging in an optically cleared coronal slice of the adult mouse brain. The use of MAP-SIM and sample-optimized illumination patterns allowed us to collect super-resolution images well beyond the typical depth limit for SIM.

## Availability of supporting data

Data is available upon request.

## Ethics approval

Because the sample was obtained commercially, animal use committee approval is not required.

## Author Contributions

TP: acquired data, analyzed data, wrote the paper.

KJ: acquired data, analyzed data.

GH: conceived project, acquired data, analyzed data, supervised research, wrote the paper.

